# Evaluation of the antioxidant activity of biologically-active compositions based on plant metabolites produced by *in vitro* cell cultures

**DOI:** 10.1101/2025.02.24.639798

**Authors:** Fedorova Anastasia Mikhailovna, Milentyeva Irina Sergeevna, Asyakina Lyudmila Konstantinovna, Prosekov Alexander Yuryevich

## Abstract

Free radicals can cause oxidative stress in the body, which, in turn, leads to various diseases such as cardiovascular diseases, diabetes, Alzheimer’s disease, Parkinson’s disease, and others. To protect against the destructive effects of radicals, living organisms utilize antioxidants. One of the most important classes of antioxidants is polyphenols. The primary effect of polyphenols on the body is associated with their ability to capture and neutralize free radicals, as well as reactive oxygen and nitrogen species. This study focuses on investigating the antioxidant activity (AA) of polyphenols and their compositions based on rutin, quercetin, and trans-cinnamic acid, produced by *in vitro* cell cultures, with the aim of developing dietary supplements with geroprotective properties. The AA of the studied substances was assessed using the ABTS and DPPH free radical scavenging methods. The study established that, according to the ABTS method, AA decreases in the order of quercetin > rutin > trans-cinnamic acid, while according to the DPPH method, the order is rutin > quercetin > trans-cinnamic acid. Among the tested mixtures, the highest AA was observed in mixture No. 8 and mixture No. 3, regardless of the assessment method. Consequently, to achieve the highest AA in a mixture, a higher content of quercetin and a minimal amount of trans-cinnamic acid should be included, while the optimal amount of rutin could not be determined. It is assumed that the amount of rutin should be at least equal to the amount of trans-cinnamic acid and ideally equal to the amount of quercetin. Thus, when developing dietary supplements with high AA, it is more advisable to include only quercetin rather than mixtures that also contain rutin and trans-cinnamic acid.

## Introduction

Currently, special attention is being given to food products that exhibit antioxidant properties, as both domestic and foreign scientists consider the excessive accumulation of free radicals to be one of the primary causes of pathological processes in the human body. This accumulation contributes to the prevalence of socially significant diseases and premature aging of the population [1].

Research conducted over the past decades has shown that free radicals can reversibly and irreversibly degrade substances belonging to all biochemical classes, including lipids and lipoproteins, nucleic acids, proteins and free amino acids, carbohydrates, and connective tissue molecules. Additionally, they negatively affect various cellular functions such as membrane activity, metabolism, and gene expression [2].

Given this, it is essential for human nutrition to include a regular intake of antioxidants – biologically-active compounds that can slow down or prevent the oxidation of organic substances, thereby protecting the body from the harmful effects of free radicals.

One approach to ensuring antioxidant protection of the body is through the use of dietary supplements (DS) containing a complex of natural antioxidants, such as polyphenolic compounds, including rutin, quercetin, and trans-cinnamic acid.

The objective of this study is to evaluate the antioxidant activity of biologically-active compositions based on plant metabolites (rutin, quercetin, and trans-cinnamic acid) produced by *in vitro* cell cultures, with the aim of developing dietary supplements with geroprotective properties.

Antioxidant activity can be assessed using various analytical methods, which are based on different properties of the studied compounds. Studies employing stable free radicals such as 2,2’-azinobis-[3-ethylbenzothiazoline-6-sulfonate] (ABTS) and 2,2’-diphenyl-1-picrylhydrazyl (DPPH) are particularly popular due to their simplicity and adaptability [3].

The proposal to use ABTS for the formation of a stable cation-radical via oxidation and for measuring the «total antioxidant capacity» was first introduced in 1993 [4]. Initially, the method was presented as an additive approach, involving the continuous formation of ABTS•+ from a substrate present in the reaction medium, catalyzed by metmyoglobin in the presence of hydrogen peroxide. The authors also used Trolox, a synthetic analog of α-tocopherol with enhanced water solubility, as a standard antioxidant. The reaction was conducted in phosphate-buffered saline (PBS) [5]. Due to potential overestimation of results caused by interactions between some antioxidants and reaction substrates or intermediate products, a modified version of the method was developed. In this improved approach, radicals were pre-generated from ABTS through a direct reaction with potassium persulfate (post-addition method) over a minimum incubation period of 6 hours. The radicals were then diluted in PBS or ethanol until an optical density of 0.7 was reached before being mixed with the sample. The antioxidant capacity was determined by measuring the remaining radicals after a predetermined reaction time (6 minutes). In their publication on the revised method, the authors demonstrated its applicability not only for food components but also for blood plasma analysis [6].

Another method for determining antiradical activity, currently the most popular, involves the deactivation of the synthetic stable radical DPPH. This approach gained widespread recognition primarily due to the work of Brand-Williams et al. [7]. The authors proposed using a methanolic radical solution (DPPH• radicals are readily available in pre-prepared form, which likely contributes to their popularity) to study the antioxidant activity of food ingredients by measuring the absorption of the remaining radicals in the reaction medium. In this and other studies [8], researchers measured the number of residual radicals until equilibrium was reached. For complete reaction with certain compounds, this process could take several hours. In later research, the procedure was simplified by setting a predefined reaction time, which typically returned to 30 minutes, as originally established in the first study using these radicals [9]. The widespread adoption of relative activity units (percentage of radical deactivation) or conversions based on the activity of a standard antioxidant – typically Trolox or ascorbic acid – has largely replaced calculations based on reaction kinetics (such as the time required for deactivation of half the radicals at a standard ratio and concentration of antioxidant and radical) [10].

### Objects and methods of research

The objects of this study were the plant metabolites rutin, quercetin, and trans-cinnamic acid, extracted from callus and root cultures *in vitro*, as well as compositions (mixtures) based on them.

Rutin was extracted from the callus culture of meadowsweet (*Filipendula ulmaria*). Quercetin was extracted from the callus culture of ginkgo biloba (*Ginkgo biloba*). Trans-cinnamic acid was extracted from the hairy root culture of Baikal skullcap (*Scutellaria baicalensis*). These metabolites, with a purity of no less than 95%, were obtained at earlier stages of research [11–15].

The composition of biologically-active substance (BAS) mixtures based on rutin, quercetin, and trans-cinnamic acid in different ratios is presented in Table 1.

**Table 1.** Composition of BAS mixtures studied in this research.

| № | Ratio of BAS in mixtures |  |  |
| --- | --- | --- | --- |
|  | Rutin | Quercetin | Trans-cinnamic acid |
| 1 | 1 | 1 | 1 |
| 2 | 2 | 1 | 1 |
| 3 | 1 | 2 | 1 |
| 4 | 1 | 1 | 2 |
| 5 | 1 | 0 | 1 |
| 6 | 1 | 0 | 3 |
| 7 | 4 | 1 | 15 |
| 8 | 0 | 3 | 1 |

For *in vitro* studies, BAS (biologically-active substances) solutions were prepared at concentrations of 1000 μM, 800 μM, 600 μM, 400 μM, and 200 μM. A stock solution of rutin (1 mg/mL) was prepared in 95% ethanol (RFK, Russia) and diluted with distilled water to the required concentrations.

The antioxidant activity (AA) of rutin, quercetin, trans-cinnamic acid, and eight mixtures based on them, at various concentrations, was assessed using the free radical scavenging methods with 2,2’-azinobis-[3-ethylbenzothiazoline-6-sulfonate] (ABTS) (Panreac/AppliChem, Spain) and 2,2’-diphenyl-1-picrylhydrazyl (DPPH) (ZHK Ecotech, Russia).

To evaluate the antioxidant activity of the tested BAS using the ABTS radical, the following solutions were prepared: stock solutions of ABTS (7 mmol/L) and potassium persulfate (Sigma-Aldrich 60489, USA) (140 mmol/L). The reaction mixture was prepared by mixing equal volumes of the two stock solutions and incubating them for 16 hours at room temperature in the dark. The final solution was diluted with ethanol until an optical density of 0.70 ± 0.10 was achieved at 734 nm. To assess antioxidant activity, 3 mL of ABTS solution was mixed with 30 μL of the test solution, incubated in the dark for 30 minutes, and then the optical density was measured at 734 nm. Results were expressed as a percentage relative to the standard ABTS solution [16]. The antioxidant activity (AA) was determined using Formula 1.

To evaluate the antioxidant activity (AA) of the studied biologically-active substances (BAS) using the DPPH free radical, a DPPH solution was prepared by dissolving 0.079 g of dry DPPH reagent (ZHK Ecotech, Russia) in 1 L of 96% ethanol. For better dissolution, the solution was placed in an ultrasonic bath (VU-09-YA-FP-04, Russia) for 15 minutes. The working (stock) solution was then diluted with 96% ethanol to achieve an optical density of 0.973 at 517 nm. For analysis 5 mL of the working DPPH solution was mixed with 50 μL of the test sample and stirred. The test tubes were incubated in complete darkness for 30 minutes. Optical density was measured at 517 nm using a ClarioStar spectrophotometer (BMG, Germany). As a control sample, the working DPPH solution was mixed with 100 μL of the solvent used for sample preparation [17]. The antioxidant activity (AA) was determined using Formula 1.

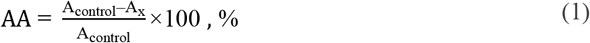

Where:

A_control_ – optical density of the working solution of ABTS or DPPH.;

A_X_ – optical density of the test solution.

## Results and discussion

During the study, the optical density of solutions containing rutin, quercetin, and trans-cinnamic acid, as well as eight mixtures based on them, was determined using a spectrophotometric method at five different concentrations: 1000 μM, 800 μM, 600 μM, 400 μM, and 200 μM. Based on the measured optical density, the antioxidant activity (AA) of the studied biologically active substances (BAS) was calculated.

The results of the AA assessment for rutin, quercetin, trans-cinnamic acid, and their eight mixtures at concentrations of 1000 μM, 800 μM, 600 μM, 400 μM, and 200 μM, measured using the ABTS free radical scavenging method, are presented in Figure 1. In this study, no control was used for AA assessment, as literature data indicate that rutin and quercetin exhibit high antioxidant activity and are commonly used as reference standards. Therefore, the AA of BAS was evaluated relative to each other, rather than against an external control.

**Figure 1.**
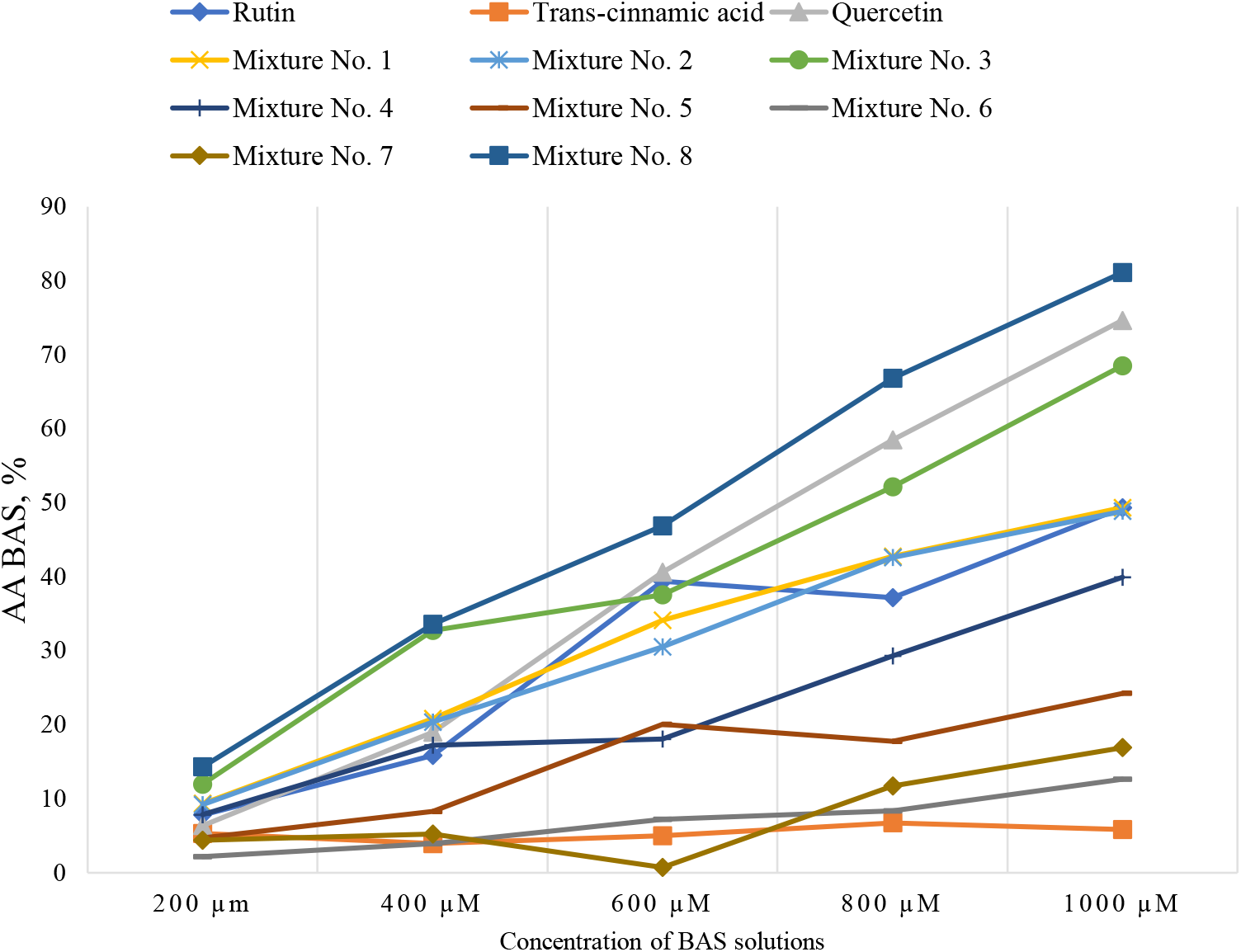
antioxidant activity of rutin, trans-cinnamic acid, quercetin, and their eight mixtures using the ABTS method

The results of the study, presented in Figure 1, indicate that the antioxidant activity (AA) of all biologically active substances (BAS) is directly proportional to the concentration of the solutions. However, an exception was observed for trans-cinnamic acid solutions at concentrations of 800 μM and 1000 μM. Among the tested solutions, quercetin demonstrated the highest AA, surpassing both rutin and trans-cinnamic acid. The maximum AA was observed for the 1000 μM quercetin solution, which exhibited 1.51 times higher AA than the rutin solution and 12.76 times higher AA than the trans-cinnamic acid solution. Conversely, the trans-cinnamic acid solution exhibited the lowest AA. These findings are consistent with previously reported literature data [18–20].

Additionally, as shown in Figure 1, the AA of BAS mixtures was also directly proportional to the concentration of the solutions, with the highest AA observed at 1000 μM. Among all tested mixtures, mixture No. 8 displayed the highest AA, which can be attributed to its high quercetin content. Excluding mixture No. 8, mixture No. 3 showed the highest AA among the remaining mixtures, again due to its relatively high quercetin content. In contrast, mixture No. 2, which contained the same amount of trans-cinnamic acid as mixture No. 3 but less quercetin and more rutin, exhibited 1.4 times lower AA than mixture No. 3. The lowest AA was recorded for mixture No. 6, which had a high proportion of trans-cinnamic acid and no quercetin. Comparing the similar AA values of mixtures No. 1 and No. 2, it was observed that the mixture with a higher quercetin content had greater activity, as determined by the ABTS method.

The results of AA assessment using the DPPH method for rutin, quercetin, trans-cinnamic acid, and their eight mixtures at concentrations of 1000 μM, 800 μM, 600 μM, 400 μM, and 200 μM are presented in Figure 2.

**Figure 2.**
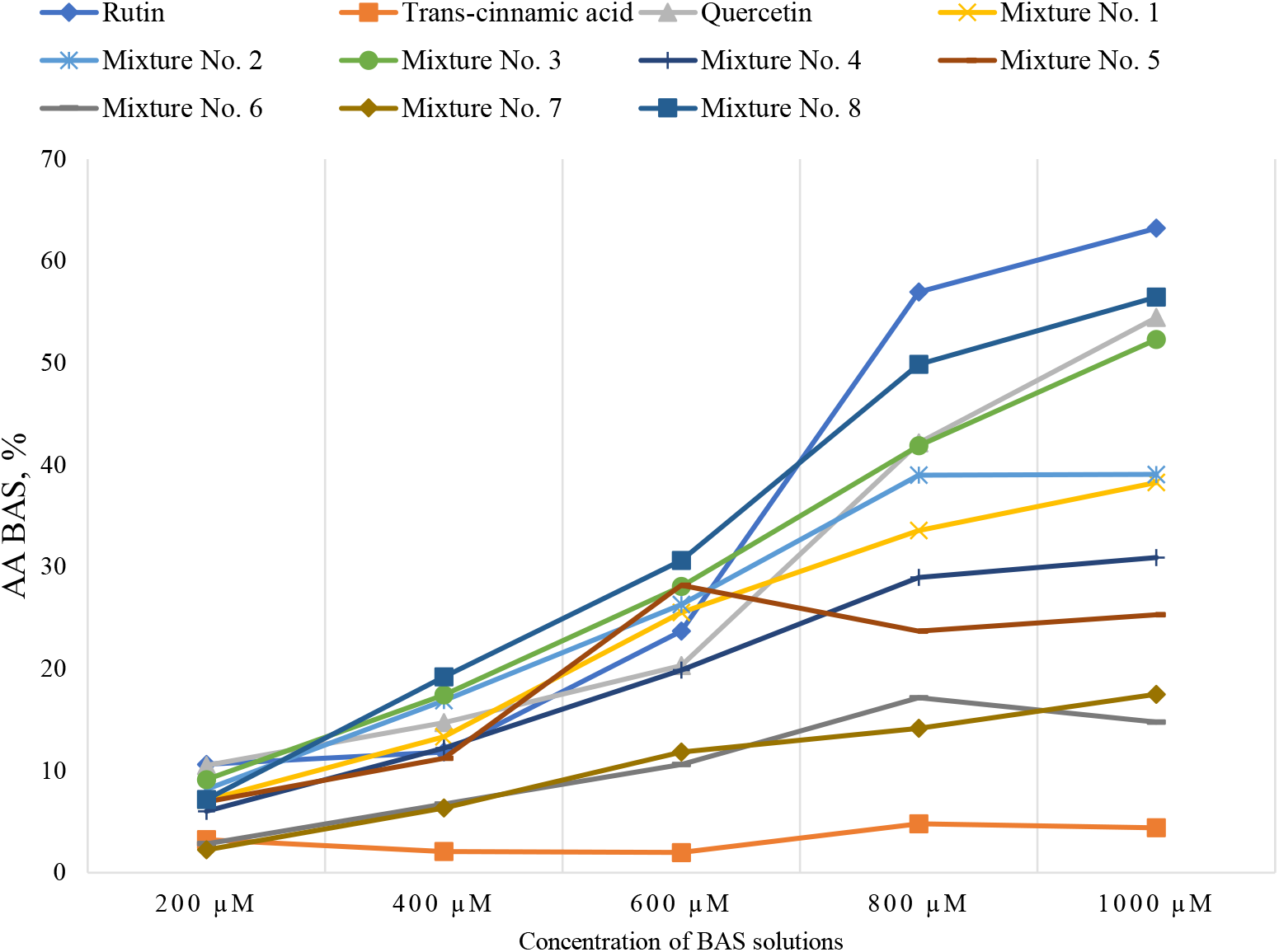
Antioxidant activity of rutin, trans-cinnamic acid, quercetin, and their eight mixtures using the DPPH method

The study revealed that the antioxidant activity (AA) of all biologically active substances (BAS) is directly proportional to their solution concentration. However, an exception was observed for the trans-cinnamic acid solution. Among the tested solutions, rutin exhibited higher AA compared to both quercetin and trans-cinnamic acid. The maximum AA was recorded for the 1000 μM rutin solution, which was 1.16 times higher than the AA of the quercetin solution and 14.3 times higher than the AA of the trans-cinnamic acid solution. The trans-cinnamic acid solution demonstrated the lowest AA. These findings are consistent with the literature data [20–22].

It was also established that the AA of BAS mixtures was directly proportional to their concentration, with the highest AA observed at 1000 μM. However, mixture No. 5 was an exception.

Despite rutin exhibiting the highest AA among individual compounds, the highest AA among the mixtures was observed for mixture No. 8, which did not contain rutin. Mixtures with a high rutin content (No. 2 and No. 5) displayed moderate AA. Excluding mixture No. 8, mixture No. 3 showed the highest AA among the remaining mixtures, likely due to its higher quercetin content. In contrast, mixture No. 2, which contained the same amount of trans-cinnamic acid as mixture No. 3 but less quercetin and more rutin, exhibited 1.33 times lower AA than mixture No. 3. The lowest AA was recorded for mixture No. 6, which had a high proportion of trans-cinnamic acid and no quercetin. Comparing the similar AA values of mixtures No. 1 and No. 2, it was observed that the DPPH method showed higher activity for the mixture containing more rutin.

It was established that mixtures containing quercetin exhibit higher antioxidant activity (AA) as determined by both ABTS and DPPH methods. Trans-cinnamic acid, regardless of the evaluation method, demonstrated weak antioxidant activity, independent of its quantity in the mixture. To achieve high AA in a mixture, the amount of rutin should not be less than that of trans-cinnamic acid and should be equal to the amount of quercetin.

However, if the goal is to develop a dietary supplement (DS) with high AA, it is more advisable to use quercetin alone, rather than mixtures that also contain rutin and trans-cinnamic acid. For example, the AA of a 1000 μM quercetin solution was 1.44 times higher than the AA of a 1000 μM solution of mixture No. 8.

## Conclusion

This study focused on evaluating the antioxidant activity (AA) of biologically active substances (BAS) and their compositions based on rutin, quercetin, and trans-cinnamic acid, produced by *in vitro* cell cultures, with the goal of developing geroprotective dietary supplements (DS). The *in vitro* AA of rutin, quercetin, trans-cinnamic acid, and eight mixtures based on them was assessed using the ABTS and DPPH methods at concentrations of 200, 400, 600, 800, and 1000 μM. The results showed that according to the ABTS method, AA decreased in the order: quercetin > rutin > trans-cinnamic acid. According to the DPPH method, AA decreased in the order: rutin > quercetin > trans-cinnamic acid. Among the mixtures, mixture No. 8 and mixture No. 3 demonstrated the highest AA, regardless of the evaluation method. Therefore, to achieve maximum AA in a mixture, it should contain a higher amount of quercetin, a minimal amount of trans-cinnamic acid, the optimal amount of rutin could not be determined. However, it is assumed that the rutin content should not be lower than that of trans-cinnamic acid and should be equal to the quercetin content. Further research is required to confirm these findings. Based on this study, it can be concluded that for the development of dietary supplements with high AA, it is more advisable to use quercetin alone, rather than mixtures containing rutin and trans-cinnamic acid.

## Funding

*This research was conducted as part of the state-funded project titled «Development of dietary supplements consisting of metabolites from in vitro plant objects for protecting the population from premature aging» (Project FZSR-2024-0008)*.

*The study was carried out using the equipment of the Shared Research Facility (SRF)*

*«Instrumental methods of analysis in applied biotechnology» at Kemerovo State University (KemSU)*.

## References

1. Kornien, N. N. Study of the antioxidant properties of food additives obtained from secondary plant resources in experiments on laboratory animals /N. N. Kornien, A.N. Troshin, A.M. Semenenko, E.V. Kuzminova [et al.] //New Technologies. – 2017. – № 1. – pp. 24–31.

2. Ludan, V. V. The role of antioxidants in the functioning of the body /V. V. Ludan, L.V. Polskaya //Taurida Medical and Biological Bulletin. – 2019. – Vol. 22, № 3. – pp. 86–92.

3. Wołosiak, R. Verification of the Conditions for Determination of Antioxidant Activity by ABTS and DPPH Assays-A Practical Approach /R. Wołosiak, B. Drużynska, D. Derewiaka, M. Piecyk and others //Molecules. – 2021. – Vol. 27(1). – P. 50. doi: 10.3390/molecules27010050/

4. Miller, N.J. A novel method for measuring antioxidant capacity and its application to monitoring the antioxidant status in premature neonates /N.J. Miller, C. Rice-Evans, M.J. Davies, V. Gopinathan, and others. //Clin Sci (Lond). – 1993. – Vol. 84(4). – pp. 407–12. doi: 10.1042/cs0840407. PMID: 8482045.

5. Miller, N. J. Spectrophotometric determination of antioxidant activity /N.J. Miller, C.A. Rice-Evans //Redox Rep. – 1996. – Vol. 2(3). – pp. 161–71. doi: 10.1080/13510002.1996.11747044.

6. Pellegrini, R. Re. Antioxidant activity applying an improved ABTS radical cation decolorization assay /R. Re, N. Pellegrini, A. Proteggente, A. Pannala, and others. //Free Radical Biology and Medicine. – 1999. – Vol. 26 (9–10). – pp. 1231–1237. 10.1016/S0891-5849(98)00315-3.

7. Brand-Williams, W. Use of a free radical method to evaluate antioxidant activity /W. Brand-Williams, M. E. Cuvelier, C. Berset. //LWT-Food science and Technology. – 1995. – T. 28. – №. 1. – pp. 25–30.

8. Bondet, V. Kinetics and mechanisms of antioxidant activity using the DPPH. free radical method /V. Bondet, W. Brand-Williams, C. Bersets Sci Technol, 2018. 7(4): p.– 1997. – T. 30. – №. 6. – pp. 609–615.

9. Blois, M. S. Antioxidant determinations by the use of a stable free radical /M.S. Blois //Nature. – 1958. – T. 181. – №. 4617. – pp. 1199–1200.

10. Kedare, S.B. Genesis and development of DPPH method of antioxidant assay /S.B. Kedare, R.P. Singh //J Food Sci Technol. – 2011. – Vol. 48(4). pp. 412–22. doi: 10.1007/s13197-011-0251-1

11. Fedorova A. M. Geroprotective activity of trans-cinnamic acid isolated from the Baikal skullcap (Scutellaria baicalensis) /A. M. Fedorova, L. S. Dyshlyuk, I.S. Milentyeva [et al.] //Food Processing: Techniques and Technology. – 2022. – Vol. 52, No. 3. – P. 582–591. – DOI 10.21603/2074-9414-2022-3-2388.

12. Faskhutdinova, E. R. Effects of bioactive substances isolated from Siberian medicinal plants on the lifespan of Caenorhabditis elegans /E. R. Faskhutdinova, A. S. Sukhikh, V.M. Le [et al.] //Foods and Raw Materials. – 2022. – Vol. 10, No. 2. – P. 340–352. – DOI 10.21603/2308-4057-2022-2-544

13. Vesnina, A. Evaluation of the In Vivo Anti-Atherosclerotic Activity of Quercetin Isolated from the Hairy Roots of Hedysarum neglectum Ledeb /A. Vesnina, I. S. Milentyeva, V. Minina [et al.] //Life. – 2023. – 13. – 1706. DOI:10.3390/life13081706

14. Faskhutdinova, E.R. Influence of Ginkgo biloba extract and its biologically-active substances on the accumulation of lipofuscin in the body of Caenorhabditis elegans. /E.R. Faskhutdinova, I.S. Milentyeva, A.I. Loseva [et al.] //Living Systems Technologies. – 2023. – Vol. 20. – № 4. – pp. 121–130. DOI: 10.18127/j20700997-202304-12.

15. Chekushkina, D. Yu. Potential use of Scutellaria baicalensis hairy root metabolite as a geroprotector /D. Yu. Chekushkina, I. S. Milentyeva, V.M. Le [et al.] //Bulletin of South Ural State University. Series: Food and Biotechnology. – 2024. – Vol. 12, № 2. – pp. 87–95.

16. Vital, A. N. S. Physicochemical composition and antioxidant activity of sweet potato flours from different cultivars produced in the Sub-middle São Francisco region /A.N.S. Vital, V.C. Benício, Y.L.F. Lins, K.W.C. Viana, C.M.B.O. Messias //Ciencia Rural. – 2023. – Vol. 53(3). – P. e20210718. 10.1590/0103-8478cr20210718

17. Maltseva, E. M. Antioxidant and antiradical activity in vitro of Sanguisorba officinalis L. extracts collected at different growth phases /E.M. Maltseva, N.O. Egorova, I.N. Egorova, R.A. Mukhamadiyarov //Medicine in Kuzbass. – 2017. – Vol. 16. – № 2. – pp. 32–38.

18. Gulya, A.P Antiproliferative and antioxidant activities of nitrate-[4-(3,4-dimethylphenyl)-2-(2-oxo-3-methoxybenzylidene)hydrazinecarbothioamide] copper /A.P. Gulya, V.S. Gudumak, O.S. Garbuz [et al.] //International Scientific Research Journal.. – 2018. – №2. – pp. 68

19. Dryakhlova, E.A. Spectrophotometric methods for determining the antioxidant activity of extracts from medicinal plant raw materials. /E.A. Dryakhlova //Current Issues of Modern Medicine and Pharmacy. – 2023.– pp. 859 – 862

20. Jo, Y. J. Antioxidant activity of β-cyclodextrin inclusion complexes containing trans-cinnamaldehyde by DPPH, ABTS and FRAP /Y.J. Jo, H.S. Cho, J.Y. Chun //Food Sci Biotechnol. – 2021. – № 30(6). – pp. 807–814. doi: 10.1007/s10068-021-00914-y.

21. Pham, T. L. Antioxidant activity of an inclusion complex between rutin and β-cyclodextrin: experimental and quantum chemical studies /T.L. Pham, T.T. Ha Nguyen, T.A. Nguyen, I. Le-Deygen, T.M. Hanh Le and others //RSC Adv. – 2024. – № 14(26). – P. 18330-18342. doi: 10.1039/d4ra02307b,

22. Ghasemi, S. Isolation and structure elucidation of the compounds from Teucrium hyrcanicum L. and the investigation of cytotoxicity, antioxidant activity, and protective effect on hydrogen peroxide-induced oxidative stress /S. Ghasemi, M. Evazalipour, N. Peyghanbari, E. Zamani and others //BMC Complement Med Ther. – 2023. – № 23(1). – P. 447. doi: 10.1186/s12906-023-04262-8.

